# Mutant fixation in the presence of a natural enemy

**DOI:** 10.1101/2023.02.10.528099

**Authors:** Dominik Wodarz, Natalia L. Komarova

**Affiliations:** Department of Population Health and Disease Prevention, University of California, Irvine, CA 92697; Department of Mathematics, University of California, Irvine, CA 92697

## Abstract

In the extensive literature about mutant invasion and fixation, populations are typically assumed to exist in isolation from their ecosystem. Yet, populations are part of ecological communities, and enemy-victim (e.g. predator-prey or pathogen-host) dynamics are particularly common. We use computational models to re-visit the established theory about mutant fixation in the presence of a natural enemy, which equally attacks both wild-type and mutant populations. We consider advantageous and disadvantageous mutants, whose fitness is unrelated to the infection. Using spatially structured agent-based as well as patch models of a population that is subject to infection, we investigate the fixation probability of a mutant that is introduced into the population at quasi-equilibrium. We find that infection significantly weakens selection. Thus, the presence of infection increases the fixation probability of disadvantageous mutants and decreases the fixation probability of advantageous mutants, with the magnitude of the effect rising with the infection rate. We show that this occurs because infection induces spatial structures, in which mutant and wild-type individuals are mostly spatially separated. Thus, instead of mutant and wild-type individuals competing with each other, it is mutant and wild-type “patches” that compete, resulting in smaller fitness differences and hence weakened selection. Because natural enemies such as infections are ubiquitous, this has broad applicability to natural systems. Our results imply that the burden of deleterious mutants in natural populations might be significantly higher than expected from mutant invasion theory developed in the absence of natural enemies.

## 1. Introduction

In reproducing populations, evolution is driven by the generation of new mutations, and the fate of the mutants is determined by selection and drift. The dynamics of mutant invasion have been studied extensively in a variety of settings [1, 2], driven in large part by the analysis of mathematical models. The fixation probability of a mutant is a central concept in this respect [1, 3, 4]. It is defined as the probability for a mutant that has been introduced into a population to rise and replace the wild-type. The conditional average time to fixation of a mutant is another important measure, determined across those instances where mutant fixation occurs. An extensive literature exists assuming constant populations [3, 5-7], which can be described mathematically by e.g. the Moran process or the Fisher-Wright process. Much of this work assumes well-mixed populations, although important insights into the dynamics of mutant invasion have been obtained for spatially or deme-structured populations [8-13], as well as more generally for mutant fixation on graphs [14-18]. Besides constant populations, the effect of demographic fluctuations around an equilibrium on the probability of mutant fixation has also been analyzed [19-24].

Evolutionary theory about mutant fixation has typically focused on the evolving population in isolation, which has given rise to many fundamental insights. In nature, however, evolving populations exist as part of an ecosystem. Natural enemies present a particularly common ecological setting. Yet, it is currently unclear how the presence of a natural enemy (that equally attacks both wild-type and mutant individuals) influences the fixation probability of a mutant. Within such a system, the evolving population can persist around an equilibrium, which at first sight seems similar to a constant population scenario. In spatially structured (and hence biologically realistic) models, however, the stable persistence of the population can be the result of continuous local extinction events coupled with migration of individuals into temporary refuge spaces without enemies, as illustrated by patch and metapopulation models [25]. Population fluctuation, frequent extinction events, and bottlenecks have been shown to change the properties of mutant invasion [20, 26, 27], and hence it is important to study the spatial dynamics of mutant invasion in the presence of a natural enemy.

Here, we study the properties of mutant invasion and fixation in spatially structured populations at quasi-equilibrium, assuming that the evolving population exists in the presence of a natural enemy. While applicable to all enemy-victim settings, the model is formulated as a population of cells that are subject to infection by a virus (regardless of cell genotype). We start by considering a spatial stochastic agent-based model and then compare its properties to those of patch models.

## 2. Results

### 2.1. Spatial agent-based model of host evolution in the presence of infection

We consider an agent-based model (ABM) on a 2D grid, where each of n_1_xn_2_ spots could be either empty or contain an uninfected or infected cell of different types (wild-type or mutant). Each time-step consists of *N*_*1*_ elementary updates, where *N*_*1*_ is the number of currently occupied sites. At each elementary update, a random cell is picked. For uninfected cells, the following events can occur. With a probability *L* the cell attempts a division. A random spot among the 8 nearest neighbors is chosen, and if unoccupied, the offspring cell is placed there. With a probability *D*, the cell dies, and with probability *1-L-D*, no event occurs. Infected cells are assumed not to divide. When selected, they can die with a probability *A*, and attempt an infection event with a probability *B*, during which a random spot is chosen among the 8 nearest neighbors. The infection proceeds if the chosen spot contains an uninfected cell. With probability 1*-A-B*, no event occurs.

In the absence of mutants, i.e. just one cell type in a spatial pathogen-host system, the dynamics have been well defined. Over time, the population sizes of uninfected and infected cells converge to a state where they fluctuate around an equilibrium (Fig 1); we will refer to the mean equilibrium size of the uninfected population as *N*_*u*_. The spatial distribution of cells, however, strongly depends on the rate of infection. For relatively low infection rates, the cells are distributed more uniformly through space (Fig 1A). For higher infection rates, however, pronounced spatial structures emerge in which moving fronts of uninfected cells are “chased” by infected cells (Fig 1B). In a particular local area, the infection drives the cell population extinct. Continuous movement of cells, through division into adjacent spots, however, leads to the persistence of the populations on a global level (across the entire grid). This recapitulates the well-known spatial refuge effect that can contribute to population persistence and spatial pattern formation in predator-prey dynamics [25].

**Figure 1:**
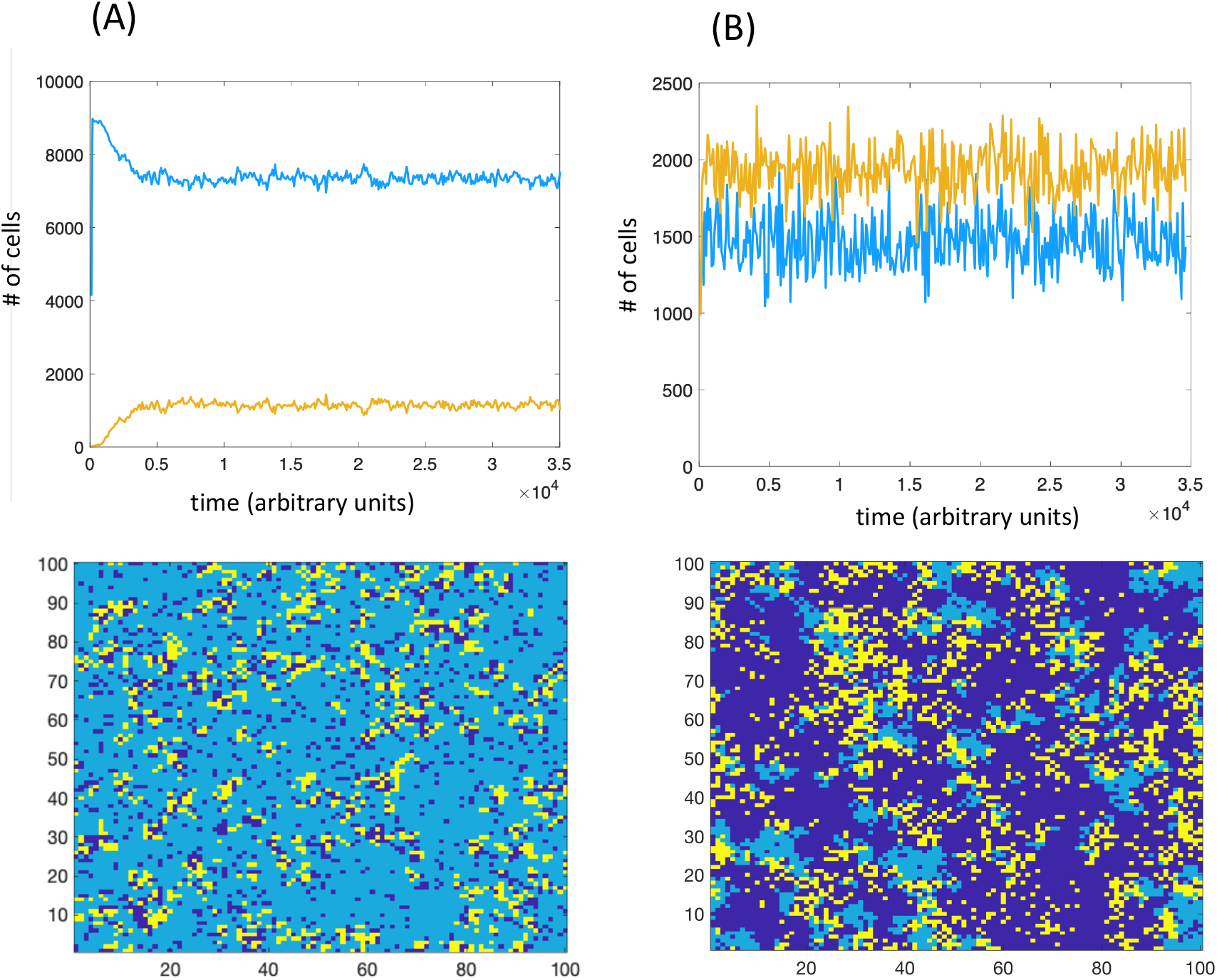
Basic dynamics of infected and uninfected cells without evolution. For the time series, light blue and yellow colors represent the populations of uninfected and infected cells. For the spatial pictures, light blue and yellow colors also represent uninfected and infected cells. Dark blue shows empty space. (A) Low infection rate. (B) High infection rate. *L=0*.*5*; *D=0*.*05*; *A=0*.*1*; *n1=n2=100*. For panel (A) *B=0*.*2*; for panel (B) *B=0*.*9*.

To study the fixation probability and conditional fixation times of mutant cells, we first simulate population dynamics in the absence of mutants for a certain amount of time, until quasi-equilibrium us reached. We then replace one (or several) randomly selected uninfected wild-type cells with a corresponding number of mutant cells, at quasi-equilibrium. This initial mutant placement occurs at a moment in the simulation when the number of uninfected wild-type cells equals the equilibrium value, *N*_*u*_, determined numerically by calculating the long-term temporal average of the population size. No *de novo* mutations are considered. We then allow the simulation to run until either the mutant is extinct, or until the mutant population has replaced the wild-type cell population (mutant fixation). The simulation is run repeatedly and the fraction of runs during which mutant fixation occurs is determined. For the cases of mutant fixation, the time until fixation is determined (conditional fixation time).

These dynamics are investigated for neutral, advantageous, and disadvantageous mutants, comparing the results in the absence and presence of infection, where the rate of infection is varied. To implement fitness advantage/disadvantage of mutants, we assume that the division rate of cells is increased/decreased relative to the wild-type by multiplying it with the coefficient (*1+s*), where *s* is the selection coefficient. An advantageous mutant corresponds to *s>0*, and for a disadvantageous mutant *s<0*.

When comparing the dynamics under different infection rates, equilibrium population sizes vary. To control for this, the grid size is adjusted such that the average equilibrium population size of uninfected cells, *N*_*u*_, is kept approximately constant, regardless of infection rate. Probabilities of mutant fixation in the presence of infection are compared with those in the absence of infection, and also with the well known formula for the Moran process of equivalent size, 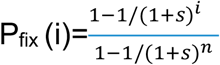, where *i* denotes the initial number of mutants in a total population of *n* individuals (*N*_*u*_ in our setting).

For neutral mutants, as expected, the numerically obtained fixation probability is 1/*N*_*u*_ regardless of infection rate. This is identical to the fixation probability given by the constant population Moran process, with the population size given by the number of uninfected cells. Only the population of uninfected cells determines the fixation probability because the infected cells are assumed to not divide in our model.

For advantageous mutants, we find that the presence of infection weakens selection (Fig 2A). Without infection, the numerically obtained mutant fixation probability is very close to the value predicted by the non-spatial Moran process (for a discussion of the role of demographic fluctuations see ref [20]); therefore, here and in other cases below, the Moran formula is a convenient reference point for evaluating the changes in fixation probability due to infection. In the presence of the virus, the fixation probability is significantly reduced, with larger reductions observed for faster infection rates. The average conditional time to fixation is found to be lower in the presence compared to the absence of infection, with shorter fixation times occurring for faster infection rates.

**Figure 2.**
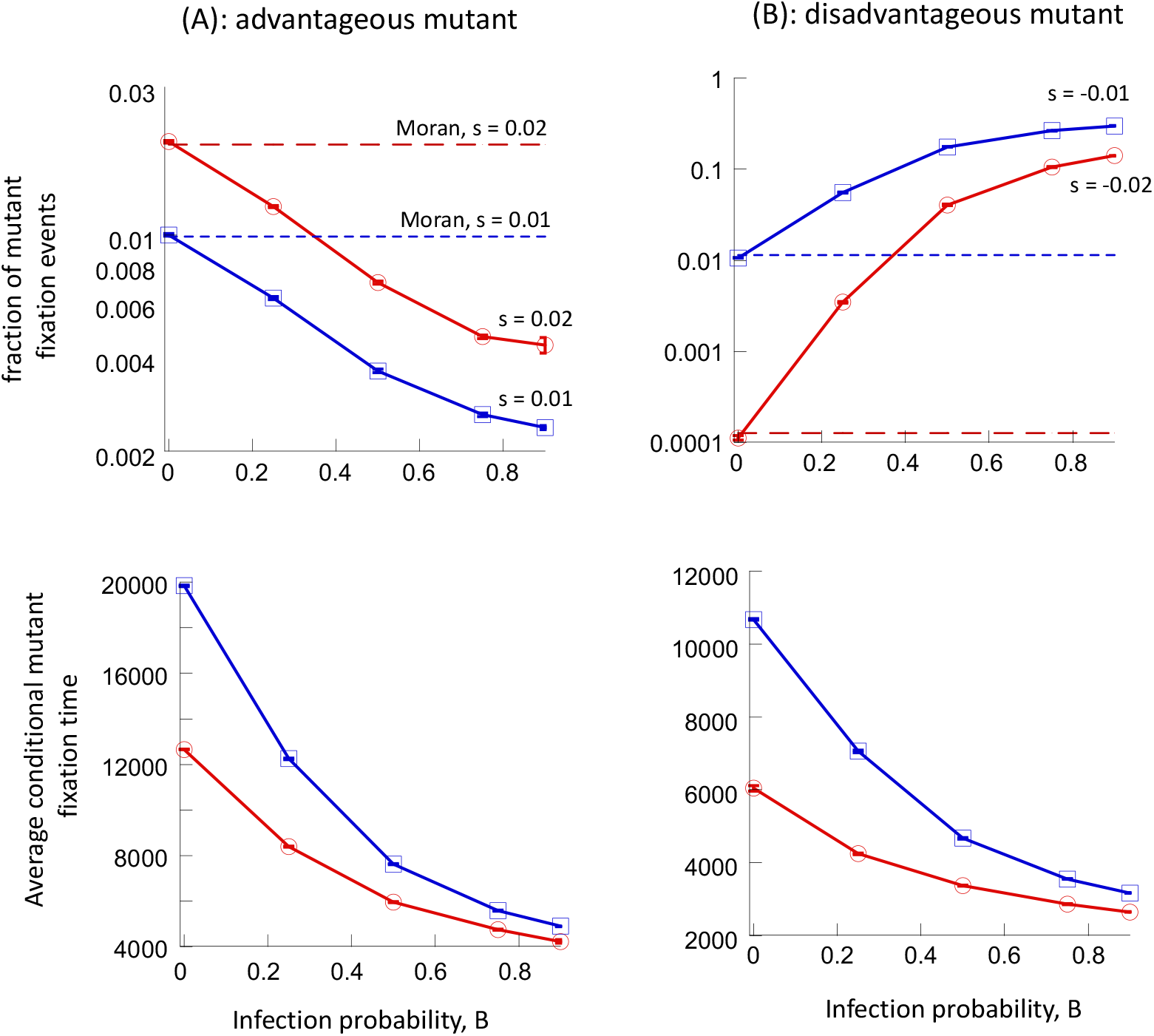
Fixation probabilities and conditional fixation times in the agent-based model for different infection probabilities. (A) Advantageous mutants for two different values of selection coefficient, s. 1 mutant was introduced. (B) Disadvantageous mutants for two different values of selection coefficient, s. 500 mutants were introduced. Parameters are as follows. *L=0*.*5*; *D=0*.*05*; *A=0*.*1*; *N*_*u*_*=944*. The grid sizes for the successive infection probabilities are: 32×33, 42×42, 60×60, 73×73, 81×80.

For disadvantageous mutants, selection is again found to be weakened in the presence of the virus (Fig 2B). Without infection, the numerically obtained mutant fixation probability is again close to the value predicted by the non-spatial Moran process. In the presence of infection, however, the fixation probability of disadvantageous mutants is significantly increased, with the larger increases seen for faster infection rates. An up to 1000-fold increase in the fixation probability is seen for the parameter regime studied here (Fig 2B). The average conditional time to fixation is again shortened by the virus (Fig 2B).

### 2.2. Deme models of host mutant evolution in the presence of infection

An alternative and coarser grained method of modeling spatial interactions are deme or patch models. Rather than tracking each individual and their spatial location, populations in the demes are assumed to be well-mixed, and their dynamics within a deme are described by stochastic Gillespie simulations of the appropriate ODEs. In addition, individuals migrate between patches, and migration can be spatially restricted to nearest neighbors, or less spatially restricted. Here we consider a two-dimensional deme / patch model, consisting of *nxn* patches. In each patch, host-pathogen dynamics are described by stochastic Gillespie simulations of ODEs, given *by dS/dt = rS(1- (S+I)/K) – δS - βSI*; *dI/dt = βSI – aI*, where *S* and *I* denote the populations of susceptible and infected cells. The parameter *r* is the basic division rate of cells, *K* is the carrying capacity of a patch, *δ* is the death rate of susceptible cells, *β* is the rate of infection, and *a* stands for the death rate of infected cells. In addition to these processes, the Gillespie simulations assume that uninfected and infected cells can migrate to a randomly chosen destination patch with a rate *m*. With spatial migration, the destination patch is chosen randomly from the eight nearest neighboring patches. For non-spatial migration, the destination patch is chosen randomly from any of the patches in the system. When varying the infection rate of the virus, the grid size is again adjusted to maintain approximately the same total number of uninfected cells across the different infection rates.

Over time, the sum of populations in this model fluctuates around a steady state value (Fig. 3). Although there is a global quasi-steady state, within each patch, populations can crash to extinction due to the infection, and can subsequently be re-colonized to repeat this pattern [25]. To investigate mutant invasion, we introduced the mutants when the total global population size of uninfected cells is equal to the rounded temporal average of the global population, *N*_*u*_. The fixation probabilities and times were determined in the same way as for the ABM.

**Figure 3.**
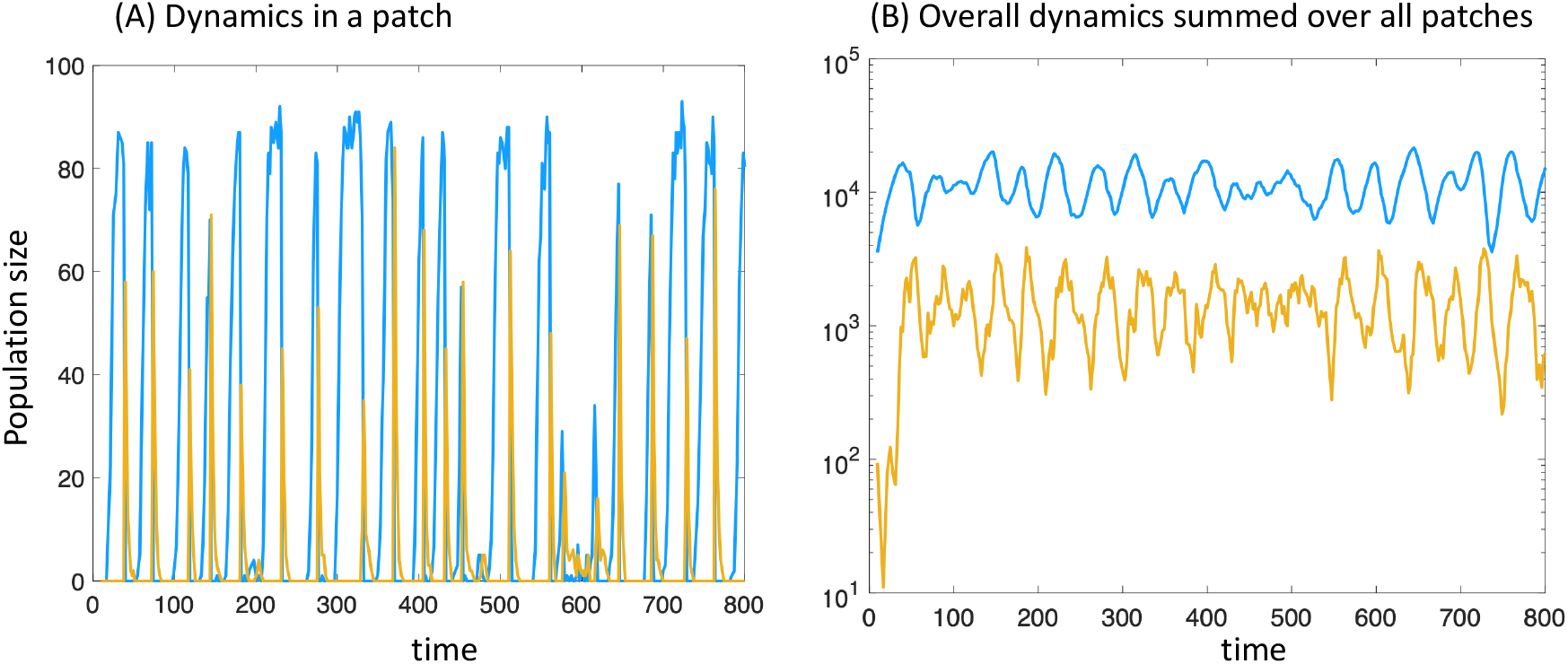
Dynamics in the patch model with migration to nearest neighboring patches. Blue and orange colors represent the populations of uninfected and infected cells. (A) Dynamics within a patch. (B) Total dynamics, with population sizes summed up across all patches. Parameters are as follows: *r=0*.*7*; *δ=0*.*1*; *β=0*.*5*; *a=0*.*5*; *K=100, m=0*.*02*, n1=n2=19.

We observe results that are qualitatively similar to the ABM described above. That is, selection is weakened both for advantageous and disadvantageous mutants (Fig 4): the fixation probability of advantageous mutants decreases below the value predicted by the non-spatial Moran process as the infection rate rises, and the fixation probability of disadvantageous mutants increases above that of the Moran process. The conditional time to fixation is generally again reduced by the presence of the infection (Fig 4), although for spatial migration the dependence can be non-monotonous. In previous work, conditional fixation times have been shown to follow complex patterns [28], and it is beyond the scope of the current study to investigate this in further detail. It is interesting to note that these trends apply both to simulations with spatial and non-spatial migration (Fig 4).

**Figure 4.**
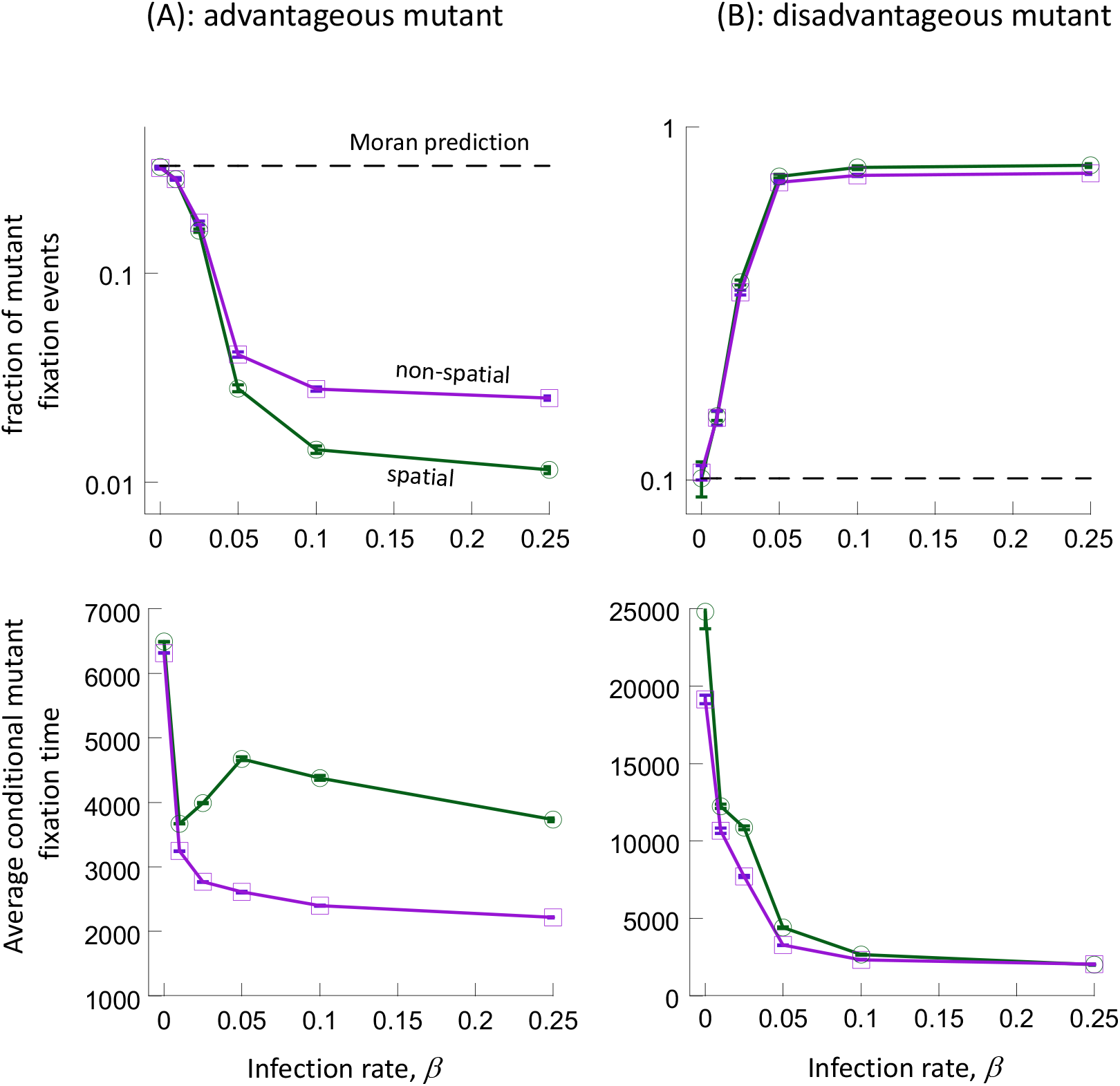
Fixation probabilities and conditional fixation times in the deme model with spatial and non-spatial migration. (A) Advantageous mutants, s = 0.02; 20 mutants were introduced. (B) Disadvantageous mutants, s = −0.001; 9000 mutants were introduced. Parameters were as follows: *r=0*.*7*; *δ=0*.*1*; *a=0*.*5*; *K=100, m=0*.*02;* Nu=11288. For spatial migration, the grid sizes for the successive infection rates are 11×12, 14×14, 22×21, 22×22, 20×21, 20×19. For non-spatial migration, they are 11×12, 14×14, 22×22, 24×24, 23×24, 22×23.

**Figure 5.**
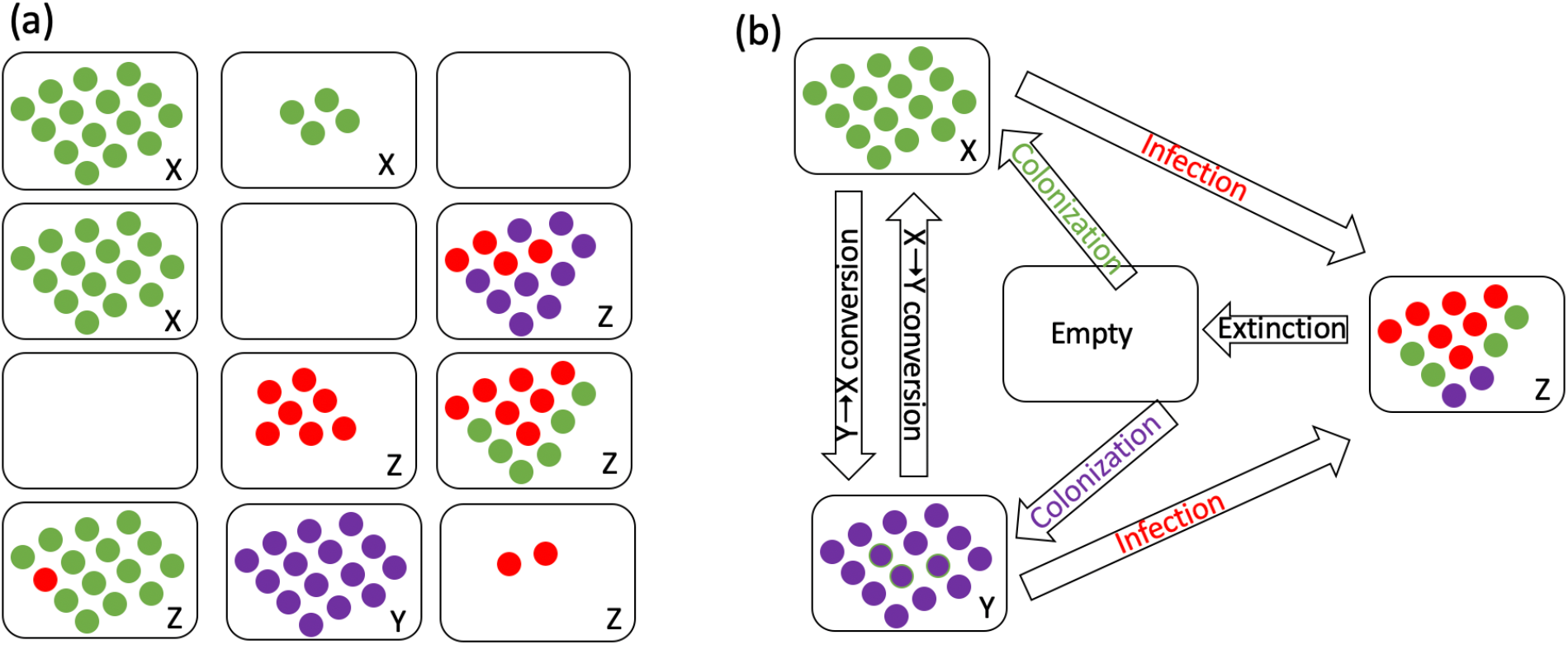
A schematic explaining the components of mutant dynamics in patches. (a) Populations of patches can be uninfected wild-type (X), uninfected mutant (Y), and infected (Z). (b) Coarse-grained model structure includes processes of colonization, infection, extinction, and conversion. Green and purple dots denote uninfected wild-type and mutant cells, and red dots indicate infected cells.

### 2.3. Patch versus cell competition

We propose that the reason for the weakened selection observed in the presence of infection is the behavior of the system as a metapopulation, regardless of the underlying model. That is, cells go extinct locally as a result of the infection, and persist by colonizing other areas of space, which temporarily do not contain infection and hence provide a refuge for the cells. This happens across a continuous space in the ABM, as shown in Fig 1. It happens more explicitly in the patch models where patches periodically go extinct and become recolonized (Fig 3), both under spatial and non-spatial migration. For relatively large infection rates, this also leads to a spatial separation of wild-type and mutant cells. In terms of the patch model, a patch is likely to either contain only wild-type cells or only mutant cells, but rarely both. In this setting, mutant and wild-type patches (rather than cells) effectively compete for colonization of empty patches, and this leads to mutant fixation probabilities that deviate significantly from those predicted by the Moran model (or the process without infection). For low infection rates or in the absence of infection, mixing of wild-type and mutant cells is more likely, and it is the competition among cells (rather than patches) that drives mutant fixation. Consequently, the observed fixation probabilities converge to those predicted by the Moran process. For intermediate infection rates, the fixation probabilities are determined by a mixture of cell and patch competition.

We demonstrate this in more detail using the patch model with non-spatial migration, see Fig.5. Assuming that mutant and wild-type cells do not co-occur in the same patches (panel (a)), we can write down a coarse-grained model where patches, rather than individual cells, are agents, and where population dynamics are governed by the following processes (panel (b)): empty patch colonization, patch infection, infected patch extinction, and patch conversion (a cell of the other type migrating into an uninfected patch and taking over). Let us denote by *X,Y*, and *Z* the total numbers of uninfected wild-type patches, uninfected mutant patches, and infected patches, and by *w*_*x*_, *w*_*y*_, and *w*_*z*_ the mean per-deme populations of these types of cells, in patches of types *X, Y*, and *Z* respectively. We can summarize the coarse-grained dynamics as follows:

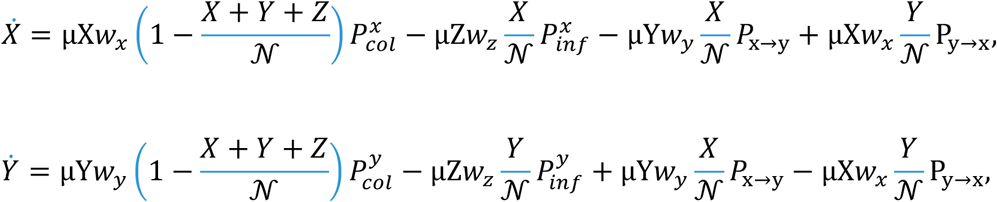

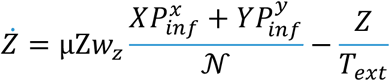

Here, the first term in the right hand side of the equations for *X* and *Y* describes colonization of empty patches, the second term describes the process of infection (the same terms reappear in the equation for *Z* with the opposite signs). The remaining terms in the *X* and *Y* equations describe conversion, and the last term in the equation for *Z* is patch extinction (see Supplement for complete details of this model). Denote the equilibrium number of uninfected patches, in the absence of mutants, by *X*_*eq*_. At the level of patch competition, there are two selection coefficients: one associated with patch conversion 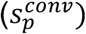 and the other with the process of extinction-recolonization 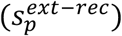. The overall selection coefficient is given by the mean of these two, weighted with their respective rates, *R*_*conv*_ and *R*_*ext-rec*_:

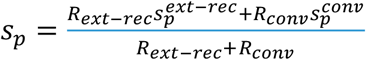, and the probability of mutant demes taking over can be calculated by the Moran formula, applied at the level of patches: 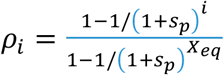, where *i* is the initial number of mutant patches. If the majority of demes are occupied and conversion is the main driving force of mutant dynamics, then the selection coefficient is approximately 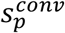, and the probability of mutant take-over is approximately that predicted by the usual Moran process in the absence of infection (and with the equivalent population size). This is because the process of deme conversion through death-birth events is the same as what comprises the conventional dynamics that leads to the Moran formula. If however the system of patches is sparsely populated and the process of recolonization is the leading force of mutant spread, then we have 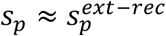, where 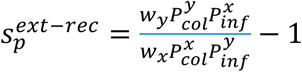, containing quantities 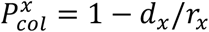 and 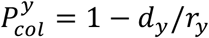, which are the probabilities for a single wild-type (mutant) cell to successfully colonize a patch, and 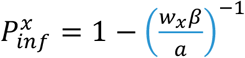 and 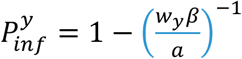, which are the probabilities for a single infected cell to start successful infection in a wild-type (mutant) patch. The selection coefficient corresponding to the process of recolonization has a significantly smaller absolute value compared to 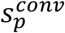, and describes much weaker selection. In the Supplement we provide a comparison with the prediction of the coarse-grained model with numerical simulations of the full system, and find them in close agreement (in the region of model applicability). While the coarse-grained approach is somewhat limited in that it does not describe patch models with spatial migration, or spatially distributed ABM systems, it provides a way to separate the different processes that contribute to mutant dynamics, and to explain the observation of a significant selection reduction in systems with infection.

## 3. Discussion

We have shown that the dynamics of mutant invasion can be significantly altered if both wild-type and mutant individuals are equally susceptible to a natural enemy such as an infection. In particular, we showed that this can lead to a significant weakening of selection. Thus, the fixation probability of a disadvantageous mutant can be strongly increased compared to the evolutionary dynamics in the absence of infection, and the fixation probability of advantageous mutants can be significantly lowered. The difference can be several orders of magnitude. In the presence of infection, it is possible that a deleterious mutant, whose invasion potential can be neglected according to traditional theory, has a reasonable chance of emerging. These results are highly relevant for natural settings, because most populations do not exist in isolation, but are part of a larger ecosystem that includes natural enemies.

We only analyzed spatially explicit models in this study, because spatial dynamics are well-known to lead to more realistic and stable enemy-victim dynamics, and lack of any spatial population structure typically results in highly oscillatory dynamics that are likely to lead to population extinction in stochastic models. Using a combination of numerical and analytic methods, we have shown that the mechanism underlying the weakened selection in this setting is a result of the population structure assumed in our models. In our system, an increase in the infection rate results in dynamics that are progressively less stable on a local level. That is, local population extinction occurs frequently, and persistence across space is possible because host populations keep moving into new locations where they can grow temporarily without the infection present [25]. In this setting, it is not the competition between individuals that matters because a given location rarely contains both mutant and wild-type individuals together. Instead, most locations contain either wild-type or mutants, and patches compete with each other in the context of extinction and recolonization of local areas / patches / demes. This leads to a much lower difference in fitness of mutant compared to wild-type individuals, accounting for the large differences seen in the fixation probabilities. These dynamics are related to extinction-recolonization dynamics that have been explored in the absence of natural enemies, also demonstrating weakened selection [26].

It is not possible to directly compare the evolutionary dynamics in the spatial systems analyzed here to a corresponding non-spatial setting. For an equivalent parameter set, the dynamics in the absence of spatial population structure result in population extinction. For other parameter regions, where the system persists in the absence of spatial structure, it is possible to investigate the fixation probability of the mutant in the presence of infection in a non-spatial system. In this case, the infection introduces an increase in demographic fluctuations around equilibrium. Previous work has shown that the existence of demographic fluctuations can independently result in weakened selection (compared to a constant population Moran process) [20]. However, this is a different mechanism, and based on preliminary simulations, selection is only weakened modestly compared to the magnitude of the effect observed in our spatial dynamics. A more detailed and comprehensive examination of the effect of an infection on demographic fluctuations and mutant fixation in non-spatial systems is subject to future work.

Experimental data that could be used to address the theoretical predictions presented here are so far not available. Bacteria-phage interactions[29] are a biological system in which these dynamics could be explored. In other contexts, bacteria and phages have been used to study aspects of virus-host evolution experimentally. Phage infections have been shown to have a direct impact on the evolution of the bacterial cell populations[30-33]. For example, the presence of bacteriophages can increase the evolutionary potential of the bacterial population, which can provide a benefit to the bacteria that balances the increased cell mortality induced by the infection[30]. In the context of antibiotic resistance evolution in bacteria, phage infections have been shown to result in the emergence of bacterial strains that are resistant to the phage, and interestingly, phage-resistance could lead to the simultaneous increase in bacterial sensitivity to the antibiotic[34, 35]. Other examples of the eco-evolutionary dynamics of phages and bacteria are discussed in references [29, 36, 37].

Our present investigation has focused on very basic evolutionary dynamics, i.e. revisiting the theory on mutant invasion if the population is subject to infection, rather than evolving in isolation. The evolutionary processes considered in our models are not in response to the selection pressure exerted by the infection, but represent infection-unrelated evolutionary changes that can be disadvantageous, advantageous, or neutral. This is in contrast to an extensive literature about host-pathogen co-evolution [38] that typically investigates scenarios where the infection is a driver of (co-)evolutionary change. The work presented here, however, has shown that the presence of an infection, or a natural enemy in general, can fundamentally alter the basic dynamics of mutant invasion in a population, through spatial heterogeneity generated by underlying dynamics.

