## Supplemental Materials for "Mutant fixation in the presence of a natural enemy"

#### Supplementary information

### 1 A coarse-grained approach in the absence of mutants

#### 1.1 The patch model of infected and uninfected cells

Consider the patch model described in the main text. We first focus on the basic dynamics in the absence of mutants. There are $\mathcal{N}$ patches, and cells migrate between patches randomly and uniformly, regardless of their location. Denote by $x_i$ and $z_i$ the populations of uninfected and infected cells in deme $i$, respectively. The co-dynamics of cells and infection within patches happen according the following ODEs:

$$\dot{x}_i = rx_i\left(1-\frac{x_i+z_i}{K}\right)-dx_i-\beta x_i z_i+\mu\left(\frac{\sum_{k=1}^{\mathcal{N}} x_k}{\mathcal{N}}-x_i\right), \quad (1)$$

$$\dot{z}_i = \beta x_i z_i - a z_i + \mu\left(\frac{\sum_{k=1}^{\mathcal{N}} z_k}{\mathcal{N}}-z_i\right), \quad 1\le i\le \mathcal{N}, \quad (2)$$

where $r$ and $d$ are respectively the linear growth and death rates of uninfected cells, $K$ is the demes' carrying capacity, $\beta$ is the infection rate, $a$ is the death rate of infected cells, and $\mu$ is the migration rate of cells. The various terms in these equations describe the propensities of all the stochastic events that are included in the Gillespie simulations used in this work.

#### 1.2 A coarse-grained description

Let us define an infected patch as a patch that contains at least one infected cell (and it may or may not contain uninfected cells). An uninfected patch contains at least one uninfected cell and no infected cells. Instead of focusing on the dynamics of infected and uninfected cells within patches, we can consider the populations of infected and uninfected patches within the grid, see figure S1

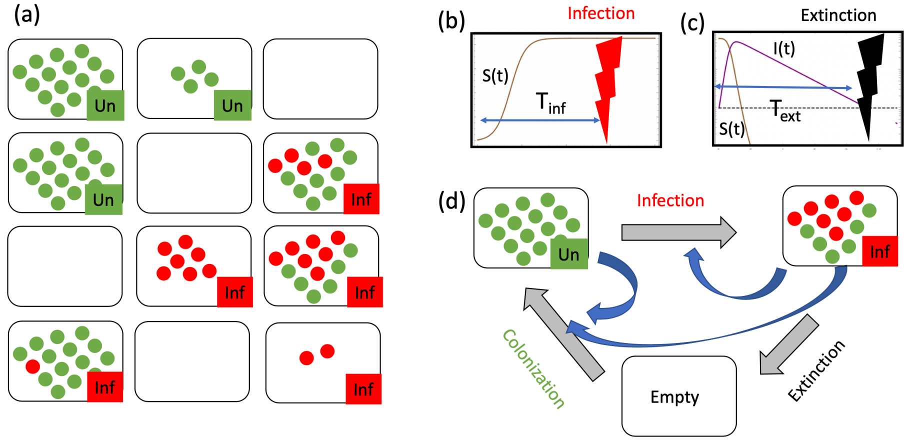

Figure S1: A schematic of the coarse-grain model. (a) Patches can be uninfected, infected, or empty; here green and red dots denote uninfected and infected cells. (b) An infected patch can become infected after a time, $T_{inf}$. (c) An infected patch can become extinct after a time, $T_{ext}$. (d) The coarse-grained description includes these two processes and a process of colonization of empty patches. Here the blue arrows indicate which populations contribute to which rates.

Suppose the number of uninfected patches is $X$, and the number of infected patches is $Z$. Let us denote the mean number of cells in an uninfected patch as $w_x^{(X)}$; the mean number of infected cells in an infected patch as $w_z^{(Z)}$, and the mean number of uninfected cells in an infected patch as $w_x^{(Z)}$. We have the following coarse-grained dynamics of patches:

$$\dot{X} = \mu(Xw_x^{(X)} + Zw_x^{(Z)})\left(1 - \frac{X+Z}{\mathcal{N}}\right)P_{col} - \mu Zw_z^{(Z)}\frac{X}{\mathcal{N}}P_{inf}, \quad (3)$$

$$\dot{Z} = \mu Zw_z^{(Z)}\frac{X}{\mathcal{N}}P_{inf} - \frac{Z}{T_{ext}}. \quad (4)$$

Here, the first term in the right hand side of equation (3) is the rate at which all the uninfected cells in the system $(Xw_x^{(X)} + Zw_x^{(Z)})$ migrate and land on an empty patch (probability to hit an empty patch is $1-(X+Z)/\mathcal{N}$), which is followed by a successful colonization (probability $P_{col}$). The second term in equation (3), as well as the first term in equation (4), describes the rate at which infected cells $(Xw_z^{(Z)})$ migrate to one of the uninfected patches (probability $X/\mathcal{N}$) and that infection takes off (probability $P_{inf}$). The last term in equation (4) is the rate at which infected patches go extinct, with $T_{ext}$ denoting the mean extinction time of an infected patch. The probability of successful colonization, $P_{col}$, and the probability of successful infection spread, $P_{inf}$, can be approximated as follows,

$$P_{col} = 1 - d/r, \quad (5)$$

$$P_{inf} = 1 - 1/R_0 = 1 - (w_x^{(X)}\beta/a)^{-1}. \quad (6)$$

Other parameters of system (4-4), such as the mean time of extinction, $P_{ext}$, and mean population sizes $w_x^{(X)}, w_z^{(Z)}$, and $w_x^{(Z)}$, are calculated from microscopic patch dynamics, as described next.

We will denote by $S(t)$ the number of uninfected cells and by $I(t)$ the number of infected cells in a typical patch.

**Uninfected patches.** We have, inside the uninfected patches:

$$\dot{S} = rS(1 - S/K) - dS - \mu S + \mu(Xw_x^{(X)} + Zw_x^{(Z)})\frac{1}{\mathcal{N}}, \quad (7)$$

$$S(0) = 1. \quad (8)$$

The life-span of an uninfected patch is defined by the timing of infection, $T_{inf}$. Infections happen at the rate $\frac{1}{\mathcal{N}}\mu Zw_z^{(Z)}P_{inf}$, so we have

$$T_{inf} = \frac{\mathcal{N}}{\mu Zw_z^{(Z)}P_{inf}}. \quad (9)$$

The mean population of an uninfected patch is therefore

$$w_x^{(X)} = \frac{1}{T_{inf}} \int_0^{T_{inf}} S(t)\, dt. \tag{10}$$

**Infected patches.** We have, inside the infected patches:

$$\begin{aligned}
\dot{S} &= rS(1-(S+I)/K) - dS - \mu S + \mu(Xw_x^{(X)} + Zw_x^{(Z)})\frac{1}{\mathcal{N}} - \beta SI, \\
\dot{I} &= \beta SI - aI - \mu I + \mu Z w_z^{(Z)} \frac{1}{\mathcal{N}}, \\
S(0) &= w_x^{(X)}, \qquad (11) \\
I(0) &= 1. \qquad (12)
\end{aligned}$$

The time to extinction, $T_{ext} > 0$, is approximated by obtaining the time that corresponds to the first minimum of the function $I(t)$. The mean population sizes are then given by

$$w_z^{(Z)} = \frac{1}{T_{ext}} \int_0^{T_{ext}} I(t)\, dt, \tag{13}$$

$$w_x^{(Z)} = \frac{1}{T_{ext}} \int_0^{T_{ext}} S(t)\, dt. \tag{14}$$

**Coarse-grained method numerical procedure.** The goal is, given all the system parameters, to predict the expected (infected and uninfected0 patch numbers and expected number of (infected and uninfected) individuals inside patches. Solutions $(X, Z)$ of system (3-4), as well as values $w_x^{(X)}, w_z^{(Z)}$, and $w_x^{(Z)}$ were used to approximate these values. The following steps were implemented:

0) Use an initial guess for the 5 quantities, $(X, Z, w_x^{(X)}, w_z^{(Z)}, w_x^{(Z)})$.

1) Evaluate parameters $P_{inf}$ and $T_{inf}$, equations (6) and (9).

2) Numerically solve the system for infected patches (equations (11-12)) up to some large value of time, $T_{max}$.

3) Obtain $T_{ext} > 0$ by locating the first minimum of the function $I(t)$ in $(0, T_{max})$.

4) Numerically solve the system for uninfected patches (equations (7-8)) up to the value $T_{ext}$.

5) Evaluate the updated values of $w_x^{(X)}$ using the solution in 4), equation (10); evaluate the updated values of $w_z^{(Z)}$ and $w_x^{(Z)}$ using the solution in 2), equations (13-14).

6) Perform an iteration of a discretized version of equations (3-4) to find updated values of $X, Z$:

$$\dot{X}^{new} = X^{old} + \Delta t \left( \mu (X^{old} w_x^{(X)} + Z^{old} w_x^{(Z)}) \left(1 - \frac{X^{old} + Z}{\mathcal{N}}\right) P_{col} - \mu Z^{old} w_z^{(Z)} \frac{X^{old}}{\mathcal{N}} P_{inf} \right), \tag{15}$$

$$\dot{Z}^{new} = Z^{old} + \Delta t \left( \mu Z^{old} w_z^{(Z)} \frac{X^{old}}{\mathcal{N}} P_{inf} - \frac{Z^{old}}{T_{ext}} \right). \tag{16}$$

7) Repeat steps (0-6), until convergence is reached.

#### 1.3 Stochastic simulations and coarse-grained predictions

Stochastic Gillespie simulations of system (1-2) were performed. The simulations were initiated by placing a relatively small block of occupied patches in the middle of a the grid, with the rest of the patches initially empty. All the occupied patches were uninfected except for a smaller block in the middle. For example, in a $40 \times 40$ grid, we started with $7 \times 7$ occupied patches in the middle, of which the central $3 \times 3$ block was infected. The initial populations of the occupied patches were just below the carrying capacity, and the initial population of infected patches contained a third of infected cells. Simulations of this type were run for a range of migration rate values, $\mu$. Figure S2 show typical dynamics inside a single patch in a multi-patch simulation, under a lower (panel (a)) and a higher (panel (b)) migration rate.

For each value of the migration rate, we obtained numerical counts of infected and uninfected patch numbers, as well as the numbers of individuals inside patches, at quasi-equilibrium. To count patch numbers, we assumed that a patch is uninfected if it contains at lest one uninfected individual and

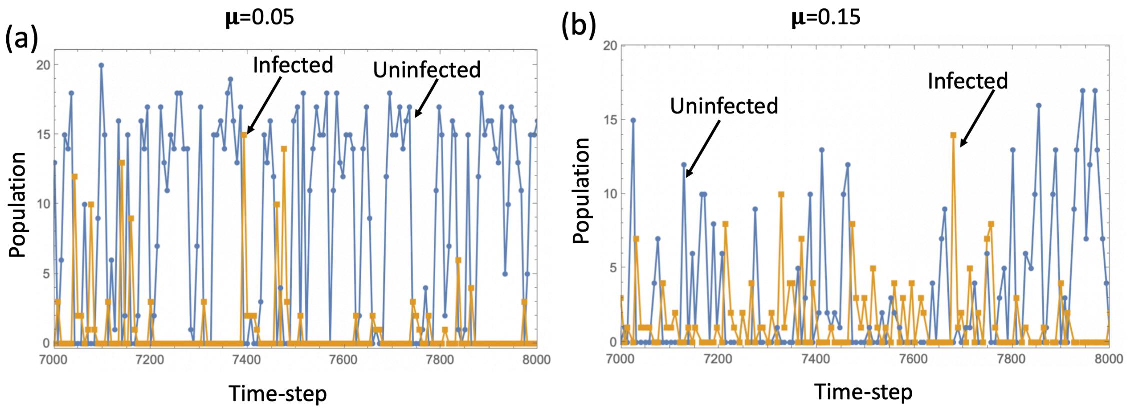

Figure S2: Typical patch trajectories under (a) low migration ($\mu = 0.05$) and (b) high migration ($\mu = 0.15$). The numbers of uninfected (blue) and infected (yellow) individuals are plotted as a function of time for a single patch. The rest of the parameters are: $r = 0.6, d = 0.1, \beta = 1.5, a = 0.5, K = 20, \mathcal{N} = 400$.

no infected individuals; a patch is infected if it contains at least one infected individual. Figure S3 shows a comparison of stochastic simulations (dots with vertical bars, representing ensemble means and standard deviations) with the coarse-grained method predictions (dashed lines), for patch numbers ($X, Z$, see panel (a)) and the numbers of individuals ($w_x^{(X)}, w_z^{(Z)}, w_x^{(Z)}$, see panel (b)).

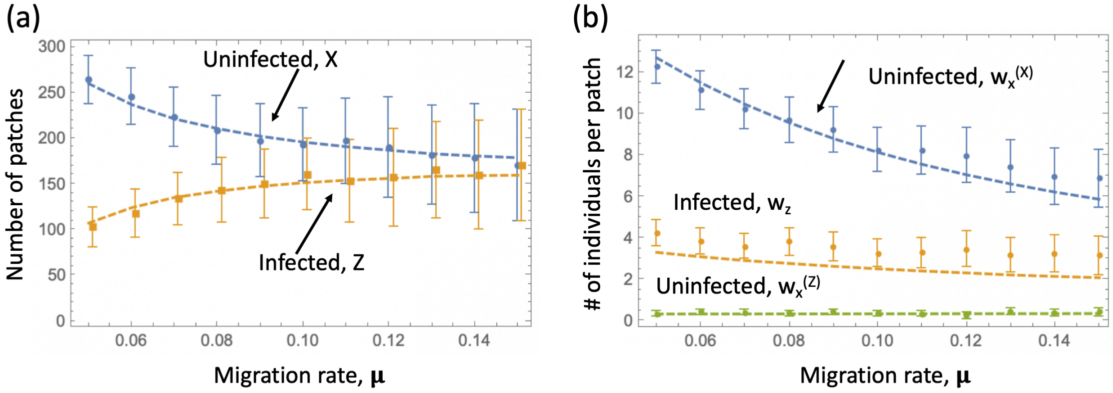

Figure S3: Comparison of stochastic simulations (dots with vertical bars, showing means and standard deviations), and the coarse-grained method predictions (dashed lines). (a) The numbers of uninfected (blue) and infected (yellow) patches are plotted for different values of the migration rate, $\mu$. (b) The numbers of uninfected individuals in uninfected patches ($w_x^{(X)}$, blue), invected individuals ($w_z^{(Z)}$, yellow), and uninfected individuals in infected patches ($w_x^{(Z)}$, green) are plotted for different values of $\mu$. The rest of the parameters are as in figure S2.

### 2 A coarse-grained approach in the presence of mutants

#### 2.1 The patch model in the presence of two types of hosts

In the presence of two types of mutants, the dynamics in the patch system are described by the following ODEs,

$$\dot{x}_i = r_x x_i \left(1 - \frac{x_i + y_i + z_i}{K}\right) - d_x x_i - \beta x_i z_i + \mu \left(\frac{\sum_{k=1}^{\mathcal{N}} x_k}{\mathcal{N}} - x_i\right), \quad (17)$$

$$\dot{y}_i = r_y y_i \left(1 - \frac{x_i + y_i + z_i}{K}\right) - d_y y_i - \beta y_i z_i + \mu \left(\frac{\sum_{k=1}^{\mathcal{N}} y_k}{\mathcal{N}} - y_i\right), \quad (18)$$

$$\dot{z}_i = \beta(x_i + y_i) z_i - a z_i + \mu \left(\frac{\sum_{k=1}^{\mathcal{N}} z_k}{\mathcal{N}} - z_i\right), \quad 1 \le i \le \mathcal{N}, \quad (19)$$

where $x_i, y_i$, and $z_i$ are respectively the number of uninfected wild-type cells, uninfected mutant cells, and infected cells. The division rates of wild-type and mutant cells are given by $r_x$ and $r_y$ respectively. The death rates of wild-type and mutant cells are given by $d_x$ and $d_y$ respectively. It is assumed that while mutants may differ from the wild type cells by their division or death rates, both types of host are equally susceptible to infection (they are characterized by the same infectivity, $\beta$), they migrate with the same rate $\mu$, and the death rate of infected cells, $a$, does not depend on their type. This is a generalization of system (1-2), and was also simulated by using the Gillespie algorithm.

#### 2.2 A coarse-grained system in the presence of two types of hosts

Let us extend the coarse-grained approach, equations (3-4), to systems with two types of host, wild type and mutant cells. In order for this approach to work, most nonempty patches in the system should contain either wild-type or mutant individuals. If a significant fraction of patches contains both types, the approach is not valid (see below for the applicability conditions). Denote by $X$ and $Y$ the numbers of uninfected wild-type and mutant patches, respectively. The numbers of wild-type and mutant infected patches are denoted respectively by $Z_x$ and $Z_y$. The coarse-grained system of equations in this case in given by

$$\begin{aligned}
\dot{X} &= -\mu(Yw_y^{(Y)} + Z_y w_y^{(Z)})\frac{X}{\mathcal{N}}P_{x\to y} + \mu(Xw_x^{(X)} + Z_x w_x^{(Z)})\frac{Y}{\mathcal{N}}P_{y\to x} \\
&+ \mu(Xw_x^{(X)} + Z_x w_x^{(Z)})\left(1 - \frac{X+Y+Z_x+Z_y}{\mathcal{N}}\right)P_{col}^x - \mu(Z_x w_z^{(Z_x)} + Z_y w_z^{(Z_y)})\frac{X}{\mathcal{N}}P_{inf}^x, \quad (20)
\end{aligned}$$

$$\begin{aligned}
\dot{Y} &= \mu(Yw_y^{(Y)} + Z_y w_y^{(Z)})\frac{X}{\mathcal{N}}P_{x\to y} - \mu(Xw_x^{(X)} + Z_x w_x^{(Z)})\frac{Y}{\mathcal{N}}P_{y\to x} \\
&+ \mu(Yw_y^{(Y)} + Z_y w_y^{(Z)})\left(1 - \frac{X+Y+Z_x+Z_y}{\mathcal{N}}\right)P_{col}^y - \mu(Z_x w_z^{(Z_x)} + Z_y w_z^{(Z_y)})\frac{Y}{\mathcal{N}}P_{inf}^y, \quad (21)
\end{aligned}$$

$$\dot{Z}_x = \mu(Z_x w_z^{(Z_x)} + Z_y w_z^{(Z_y)})\frac{X}{\mathcal{N}}P_{inf}^x - \frac{Z_x}{T_{ext}^x}, \quad (22)$$

$$\dot{Z}_y = \mu(Z_x w_z^{(Z_x)} + Z_y w_z^{(Z_y)})\frac{Y}{\mathcal{N}}P_{inf}^y - \frac{Z_y}{T_{ext}^y}. \quad (23)$$

As before, the populations of individuals in different patches are denoted with $w$, where the subscript refers to the population type ($x/y$ for uninfected w.t./mutants, and $z$ for infected cells), and the superscript refers to the patch type.

Compared to system (3-4), the equations in the presence of mutants have two new terms. The first term in the right hand side of equations (20) and (21) describes the rate at which uninfected mutant cells (totaling $Yw_y^{(Y)} + Z_y w_y^{(Z)}$) migrate to an uninfected wild-type patch (probability $X/\mathcal{N}$), and subsequently, the patch is "converted" to becoming a mutant patch by means of stochastic fixation of mutants (probability $P_{x\to y}$). Similarly, the second term in the right hand side of equations (20) and (21) describes the conversion of mutant patches to wild-type patches.

In addition, in the presence of two types of hosts, some of the rates now have a superscript that specifies the type of a patch where the process takes place. For example, $P_{col}^x$ and $P_{col}^y$ denote the probability of successful colonization of an empty patch by wild type and mutant cells, respectively, and are given by $P_{col}^x = 1 - d_x/r_x$, $P_{col}^y = 1 - d_y/r_y$.

#### 2.3 Quantification of selection dynamics

The total number of uninfected wild-type cells in system (20-23) is given by $Xw_x^{(X)} + Z_x w_x^{(Z)}$, and the total number of uninfected mutants by $Yw_y^{(Y)} + Z_y w_y^{(Z)}$. The two contributions are unequal, with the large majority of uninfected cells coming from uninfected patches. Therefore, we can ignore[1] the terms $Z_x w_x^{(Z)}, Z_y w_y^{(Z)}$ compared to $Xw_x^{(Z)}, Yw_y^{(Z)}$. The resulting 3-equation system is presented in the main text, where we denoted $Z = Z_x + Z_y$.

We would like to determine the selection pressure experienced by the mutant patches, given that they are at low numbers compared to the number of wild-type patches. To proceed, we will find the equilibrium solution of system (20-23) in the absence of mutants ($X = X_{eq}, Y = 0, Z_x = Z_{eq}, Z_y = 0$), and then linearize the system around this solution, to obtain the equation for $\dot{Y}$.

[1]Note that this step significantly simplifies all the expressions, but it is not necessary for the method to work. If we keep contributions $Z_x w_x^{(Z)}, Z_y w_y^{(Z)}$, then the equilibrium value $Z_{eq}$ can be obtained from a quadratic, rather than linear, equation, and the system for mutants will contain linear ODEs for $\dot{Y}$ and $\dot{Z}_y$, where the latter equation can be solved in quasi-equilibrium, reducing the system to a single equation for $\dot{Y}$.

We have the following equilibrium solution in the absence of mutants:

$$X_{eq} = \frac{\mathcal{N}}{\mu P^x_{inf} T_{ext} w_z^{(Z)}}, \quad Z_{eq} = \frac{\mathcal{N} w_x^{(X)} P^x_{col}}{\mu w_z^{(Z)} P^x_{inf} T_{ext}} \frac{\mu w_z^{(Z)} P^x_{inf} T_{ext} - 1}{w_x^{(X)} P^x_{col} + w_z^{(Z)} P^x_{inf}}.$$

Linearizing equation (21), we obtain

$$\dot{Y} = (\nu_{col} + \nu_{x\to y} - \nu_{inf} - \nu_{y\to x})Y, \tag{24}$$

where the four rates on the right hand side correspond to the four processes that can change the number of mutant patches:

- Colonization of empty patches by mutant cells, which increases the number of mutant patches with the per-patch rate

$$\nu_{col} = \mu w_y^{(Y)} \left(1 - \frac{X_{eq} + Z_{eq}}{\mathcal{N}}\right) P^y_{col},$$

- Conversion of wild-type patches to mutant patches, which also increases the number of mutant patches, with the per-patch rate

$$\nu_{x\to y} = \mu w_y^{(Y)} \frac{X_{eq}}{\mathcal{N}} P_{x\to y},$$

- Infection of mutant patches which leads to their subsequent extinction (and decreases their number) at the per-patch rate

$$\nu_{inf} = \mu Z_{eq} w_z^{(Z)} \frac{1}{\mathcal{N}} P^y_{inf},$$

- Conversion of mutant patches to wild-type patches, which decreases the number of mutant patches with the per-patch rate

$$\nu_{y\to x} = \mu X_{eq} w_x^{(X)} \frac{1}{\mathcal{N}} P_{y\to x}.$$

To better understand the role of these processes in patch selection, we can envisage a corresponding coarse-grained stochastic process, where the number of mutant patches is denoted as $j$, and during an infinitesimal time, $\Delta t$, the following processes take place:

- The number of mutant patches can increase by one with probability $P(j \to j+1) = j(\nu_{col} + \nu_{x\to y})\Delta t$;
- The number of mutant patches can decrease by one with probability $P(j \to j-1) = j(\nu_{inf} + \nu_{y\to x})\Delta t$;
- The number of mutant patches can remain constant with probability $P(j \to j) = 1 - (P(j \to j+1) + P(j \to j-1))$.

Denote by $\rho_j$ the probability that starting from $j$ mutant patches, the system reaches the state where all patches are mutant. We have

$$\rho_j = P(j \to j+1)\rho_{j+1} + P(j \to j-1)\rho_{j-1} + \rho_j(1-(P(j \to j+1)+P(j \to j-1))),$$

which is equivalent to

$$\rho_{j+1}(\nu_{col} + \nu_{x\to y}) + \rho_{j-1}(\nu_{inf} + \nu_{y\to x}) - \rho_j(\nu_{col} + \nu_{inf} + \nu_{x\to y} + \nu_{y\to x}) = 0.$$

The probability of mutant fixation is then given by

$$\rho_i = \frac{1 - 1/(1+s_p)^i}{1 - 1/(1+s_p)^N},$$

where $N \approx X_{eq}$ and the effective patch selection coefficient, $s_p$, is given by

$$s_p = \frac{\nu_{col} + \nu_{x\to y}}{\nu_{inf} + \nu_{y\to x}} - 1 = \frac{R_{ext-rec} s_p^{ext-rec} + R_{conv} s_p^{conv}}{R_{ext-rec} + R_{conv}},$$

where the rates of the two processes are

$$R_{conv} = \frac{X_{eq}}{\mathcal{N}} \mu w_x^{(X)} P_{y\to x} = \frac{w_x^{(X)} P_{y\to x}}{w_z^{(Z)} P_{inf}^x T_{ext}}, \tag{25}$$

$$R_{ext-rec} = \frac{Z_{eq}}{\mathcal{N}} \mu w_z^{(Z)} P_{inf}^y = \frac{P_{inf}^y w_x^{(X)} P_{col}^x (\mu w_z^{(Z)} P_{inf}^x T_{ext} - 1)}{P_{inf}^x T_{ext}(w_x^{(X)} P_{col}^x + w_z^{(Z)} P_{inf}^x)}, \tag{26}$$

and the associated effective selection coefficients are

$$s_e^{conv} = \frac{w_y^{(Y)}}{w_x^{(X)}} \frac{P_{x\to y}}{P_{y\to x}} - 1, \quad s_e^{ext-rec} = \frac{w_y^{(Y)}}{w_x^{(X)}} \frac{P_{col}^y}{P_{col}^x} \frac{P_{inf}^x}{P_{inf}^y} - 1.$$

The regime where the extinction-recolonization process is dominant is characterized by

$$Z_{eq} w_z^{(Z)} P_{inf}^y \gg X_{eq} w_x^{(X)} P_{y \to x},$$

or, denoting by $N_{inf}$ and $N_{uninf}$ the total expected number of infected and uninfected cells at the mutant-free equilibrium, we have the condition

$$N_{inf} P_{inf}^y \gg N_{uninf} P_{y \to x}.$$

If the process of extinction-recolonization is predominant, then the probability of mutant fixation, starting from a single fully mutant deme, is given by

$$\rho_{patch} = \frac{1 - 1/(1 + s_e^{ext-rec})}{1 - 1/(1 + s_e^{ext-rec})^{X_{eq}}}. \quad (27)$$

#### 2.4 Comparison with simulations

If the process of conversion is dominant, then we expect the probability of mutant fixation to be similar to that of the Moran process. On the other hand, if the extinction-recolonization process is predominant compared to the conversion process, we expect selection to be significantly weaker and the probability of fixation much larger (smaller) for disadvantageous (advantageous) mutants.

To test this theory, it is instructive to use the parameter regime where the process of extinction-recolonization process is predominant. This is observed when the mutant and wild type cells are typically separated, that is, they do not co-occur in the same demes. Such separation happens for example if many demes are unoccupied (allowing for recolonization by a single type), and when demes' life-span is relatively short (due to intense infection), which precludes accumulation of cells of the different types due to migration.

While spatial separation of mutants and wild-types is typical in simulation with spatially restricted migration, it is more difficult to achieve under non-spatial migration. To increase the degree of separation of mutants and wild types (and thus to amplify the contribution of extinction-recolonization process in the absence of space), we assumed that infected cells migrated at a faster rate than uninfected cells. As a result, demes' lifespans became shorter (due to intense infection) and conversion did not play a significant role.

Figure S4 shows a system where mutant and wild-type cells are largely separated. In a simulation run where mutant individuals expand from low

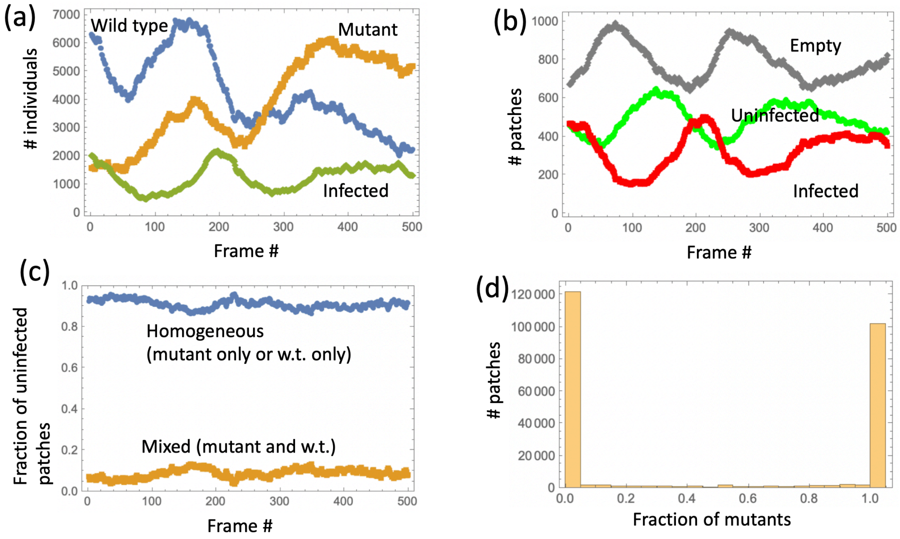

Figure S4: Dynamics of a patch system with non-spatial migration. (a) The total number of uninfected wild type (blue) and mutant (orange) individuals, as well as infected (green) individuals, as a function of frame number. (b) The number of uninfected (green), infected (red), and empty (gray) patches, as a function of frame number. (c) the fraction of uninfected patches (that is, patches that only contain either mutant or wild type cells), blue, and mixed (both mutant and wild type cells, orange) patches, as a function of frame number. (d) A histogram showing the number of uninfected nonempty patches characterized by different fractions of mutant cells – a congregate over all 500 frames. The parameters are $r = 0.7, d = 0.6, K = 200, a = 0.1, \beta = 0.04, N = 1600$. The migration rate is $\mu = 0.01$ for uninfected individuals and twice that amount for infected individuals.

numbers to reach a majority, we recorded the state of the patches after every 10,000 updates, creating a total of 500 frames. Here, the state of the patches is described by the number of uninfected wild type, uninfected mutants, and infected individuals. Panel (a) shows a time-series for the total (over all patches) numbers of these three types of individuals, to show that the overall number of mutants varies from low to high numbers. Panel (b) shows three types of patches: uninfected (that is, patches that only contain uninfected individual, regardless of their type), infected (that is, patches that contain at least one infected individual), and empty. In particular, we observe that the number of empty patches is relatively high (the total number of patches is 1600). Panel (c) plots time-series that characterize the degree of mixing of wild type and mutant individuals: for all uninfected nonempty patches, we plot the fraction of patches that either contain mutant or wild type cells (the homogeneous patches, blue); similarly, in orange, we plot the fraction of mixed patches (that is uninfected patches that contain at least one wild-type and at least one mutant individual). We observe that the fraction of mixed patches stays low throughout the frames. Finally, panel (d) shows a histogram of mutant fraction in the uninfected patches over all 500 frames; we can see again that the majority of patches are either 100% mutant or 100% wild-type.

Note that most simulations that start with a low number of mutants will result in mutant extinction without ever seeing a significant number of mutants; for such runs, it is not surprising that most of the patches are homogeneous (and contain wild-type cells). The simulation shown in figure S4 was chosen such that mutants do rise to a significant percentage. It demonstrates that even in such cases, the two types of individuals remain mostly separated (that is, rarely inhabit the same patch).

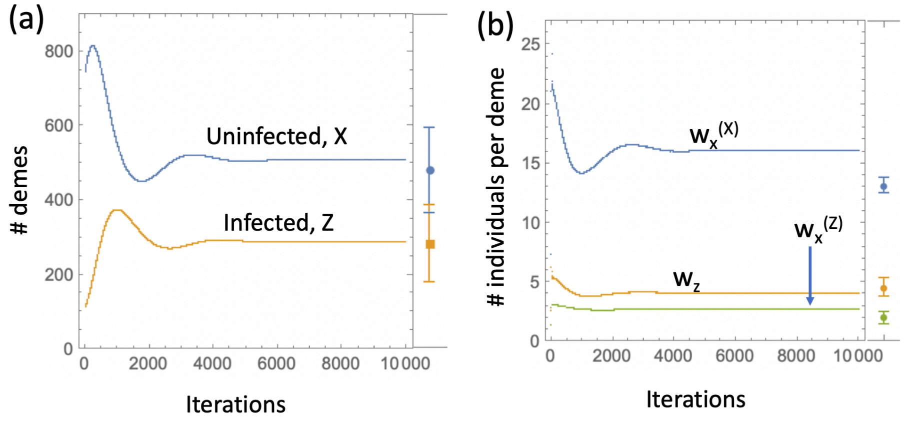

Figure S5: The coarse-grained approach applied to the system in figure S4. Iterations of steps (0-6) of section 1.2 are shown for the deme numbers (a) and population sizes (b). The means and standard deviations obtained by stochastic simulations are shown on the right side of each graph. Parameters are as in figure S4.

For this system, we demonstrate that the method described above produces a very good prediction of the mutant fixation probability. Figure S5 shows the result of iterations of system (15-16), to obtain an approximation of the deme numbers and population sizes. The means and standard deviations obtained by stochastic simulations are shown on the right side of each graph.

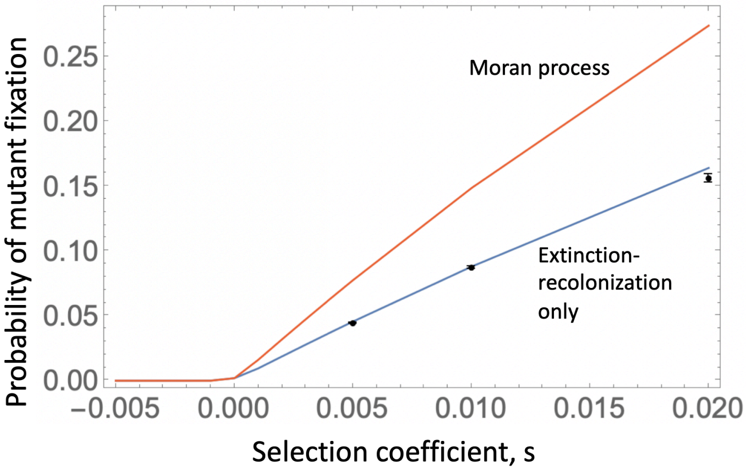

Figure S6: Mutant fixation probability as a function of mutant selection coefficient, $s$. The blue line is the coarse-grained prediction, equation (27). The red line is the Moran prediction, equation (28). The points with small vertical bars represent the mean and standard deviation of the probability of fixation determined by stochastic simulations for three different values of $s$. Parameters are as in figure S4.

The probability of mutant fixation, starting with a single fully-mutant deme, is given by equation (27). Figure S6 compares this calculation (blue line) with the result for the usual Moran prediction,

$$\rho_{Moran} = \frac{1 - 1/(1+s)^{w_x^{(X)}}}{1 - 1/(1+s)^{w_x^{(X)} X_{eq}}}, \tag{28}$$

which is shown by the red line. Numerical simulations of the stochastic system were performed to estimate the probability of mutant fixation; the results are shown by points with small standard deviations that correspond to three different values of the mutant selection coefficient $s$. In these simulations, first, a patch system in the absence of mutants was allowed to reach a quasi-equilibrium. Then, at a time when the number of uninfected cells was given by $N_u$ (the numerically determined mean number at quasi-equilibrium), a patch that contained at least one uninfected cell was selected randomly, and all uninfected cells were replaced by mutant uninfected cells. The simulation was stopped when either the mutants reached fixation or were extinct. This process was repeated $15,480$ times, for each value of $s$, yielding the fraction of the runs that resulted in mutant fixation.

We observe that the prediction of the coarse-grained approach is very similar to the observed probability of mutant fixation, and that the Moran prediction corresponds to a much higher fixation probability for advantageous mutants.
